# Axon initial segment plasticity caused by auditory deprivation degrades time difference sensitivity in a model of neural responses to cochlear implants

**DOI:** 10.1101/2024.12.10.627765

**Authors:** Anna Jing, Sylvia Xi, Ivan Fransazov, Joshua H. Goldwyn

**Affiliations:** Swarthmore College, Department of Mathematics and Statistics. 500 College Ave. Swarthmore, PA, 19081, USA

**Keywords:** cochlear implant, axon initial segment, plasticity, medial superior olive, sound localization, computer model

## Abstract

Synaptic and neural properties can change during periods of auditory deprivation. These changes may disrupt the computations that neurons perform. In the brainstem of chickens, auditory deprivation can lead to changes in the size and biophysics of the axon initial segment (AIS) of neurons in the sound source localization circuit. This is the phenomenon of axon initial segment (AIS) plasticity. Individuals who use cochlear implants (CIs) experience periods of hearing loss, and so we ask whether AIS plasticity in neurons of the medial superior olive (MSO), a key stage of sound location processing, would impact time difference sensitivity in the scenario of hearing with cochlear implants. The biophysical changes that we implement in our model of AIS plasticity include enlargement of the AIS and replacement of low-threshold Potassium conductance with the more slowly-activated M-type Potassium conductance. AIS plasticity has been observed to have a homeostatic effect with respect to excitability. In our model, AIS plasticity has the additional effect of converting MSO neurons from phasic firing type to tonic firing type. Phasic firing is known to have greater temporal sensitivity to coincident inputs. Consistent with this, we find AIS plasticity degrades time difference sensitivity in the auditory deprived MSO neuron model across a range of stimulus parameters. Our study illustrates a possible mechanism of cellular plasticity in a non-peripheral stage of neural processing that could impose barriers to sound source localization by bilateral cochlear implant users.

## 1 Introduction

Cochlear implants (CIs) are neural prostheses that can produce a sense of auditory perception by stimulating the auditory nerve (AN) with electrical signals. These devices can improve users’ speech understanding (Krueger et al., 2008; Gaylor et al., 2013, e.g.) and improve perception of other important aspects of sound (Wilson and Dorman, 2008, for review). For some individuals, CIs are implanted in both ears (bilateral CIs). The benefits of implanting a second CI include improved speech understanding in noise and improved sound source localization relative to users of a single CI (unilateral implant) (van Hoesel, 2004; Brown and Balkany, 2007). Compared to normal hearing listeners, however, users of bilateral CIs exhibit decreased sound source localization accuracy (Grantham et al., 2007).

Two primary cues for sound source location that normal hearing listeners use are the difference in sound level at the two ears (interaural level difference, ILD) and the difference in the arrival times of sounds at the two ears (interaural time difference, ITD). Bilateral CI users primarily rely on ILDs to determine sound source location (van Hoesel and Tyler, 2003; Seeber and Fastl, 2008). Factors that may limit the utility of ITD cues for bilateral CI users include mismatched electrode positions in the two ears (Kan et al., 2013), differences in AN temporal responses in response to CI and acoustic stimuli, and AN fiber degeneration that can vary depending on the etiology and duration of deafness. A number of modeling studies have explored how neuron death, demylenation, or other degenerative effects may impact AN activation by CIs (Briaire and Frijns, 2006; Resnick et al., 2018; Goldwyn et al., 2010), but none (to our knowledge) have considered how neural degeneration impacts sound source localization by CI users.

Periods of hearing loss can induce degenerative and plastic changes in neurons across all stages of the auditory nervous system. In this work, we focus on changes that can occur at the site of spike initiation in auditory neurons due to periods of auditory deprivation. Specifically, we draw on a number of studies that demonstrate changes to the spike initiation regions of various auditory neurons during periods of auditory deprivation (Kuba et al., 2010; Kuba, 2012; Kuba et al., 2014, 2015; Kim et al., 2019). This phenomenon is known as axon initial segment (AIS) plasticity (Yamada and Kuba, 2016). AIS plasticity is conceptualized as a homeostatic response to changed synaptic inputs (Yamada and Kuba, 2016). During periods of decreased excitatory input rates to a neuron (by ablation of the cochlea, in experiments), the neuron responds by increasing its intrinsic excitability, thereby compensating for the diminished excitatory input. Changes to the neuron during AIS plasticity include enlarged AIS size, and alterations to both the density and composition of a variety of potassium channels.

We investigate how AIS plasticity may impact the dynamics of neurons in the medial superior olive (MSO). MSO neurons are a critical component of mammalian sound-localization circuit that encodes ITDs. MSO neurons function as temporallyprecise coincidence detectors that encode small time differences in sound arrival times at the two ears (Grothe et al., 2010; McAlpine and Grothe, 2003, for review). Although AIS plasticity has not been observed in human MSO neurons, we consider AIS plasticity in these neurons plausible in the context of human CI users. We believe this because CI users experience prolonged periods of hearing loss, during which time the inputs to their auditory systems are diminshed. Additionally, binaural neurons in the ITD-detecting circuit of chickens (*nucleus magnocellularis* neurons, NM) exhibit AIS plasticity during auditory deprivation (Kuba et al., 2014) and binaural coincident detector neurons in chickens in the *nucleus laminaris* (NL) region (the homologue of MSO) have AIS regions with lengths tuned to the frequency of their synaptic inputs (Kuba et al., 2006; Kuba, 2012).

We hypothesized that if there is AIS plasticity in MSO neurons in a manner similar to what has been observed in NM neurons, then MSO neurons would become over-excitable and less able to perform their role as temporally-precise coincidence detectors of well-timed binaural excitatory inputs. We tested this hypothesis by simulating MSO neural activity driven by auditory nerve responses to bilateral CI input over a range ITD values. We compared responses of the model of MSO neurons with standard parameter values (Rothman and Manis, 2003; Khurana et al., 2011) to responses of a model with biophysical parameters adjusted to reflect AIS plasticity caused by auditory deprivation.

In our previous studies, we showed that temporally-precise coincidence detection by auditory brainstem neurons depends on optimally calibrated soma and axon regions (Goldwyn et al., 2019; Drucker and Goldwyn, 2023). Consistent with those studies, we show here that alteration to the soma-axon structure by AIS plasticity degrades ITD sensitivity. In particular we find that, although AIS plasticity is thought of as a homeostatic change with respect to firing rate, the nature of the neural dynamics (the “firing type”) changes in a consequential way. In our hands, AIS plasticity converts the model from phasic firing type in the control parameter setting to tonic firing type after AIS plasticity. Principal cells of the MSO are phasic (Svirskis et al., 2002), as are the homologous ITD detectors in the nucleus laminaris of birds (Reyes et al., 1996). Phasic dynamics enhance ITD sensitivity in MSO and NL neurons (Meng et al., 2012; Huguet et al., 2017; Drucker and Goldwyn, 2023).Accordingly, we find a loss of time-difference sensitivity when AIS plasticity converts the model to tonic type from phasic type.

In Methods we describe our implementation of the model for AN spiking in response to CI stimulation. Our model is an implementation of an established model for AN responses to CI stimulation (Boulet, 2016; Tabibi et al., 2021). We also describe the MSO model and the changes we make to the AIS based on the observations of AIS plasticity by Kuba and colleagues (Kuba et al., 2010; Kuba, 2012; Kuba et al., 2014, 2015). In Results we present how AIS plasticity degrades time-difference processing in MSO neurons in this case of responses to CI stimulation. We present our main results in terms of time-difference tuning curves (firing rates as a function of the difference in the timing of two streams of pulses that represent the CI stimulation at two different ears). We close in the Discussion section by commenting on the implications of our results for ITD-based sound localization by bilateral CI users.

## 2 Methods

In this section, we describe the two stages of our computational model. In the first stage, pulse trains of current evoke spikes in auditory nerve (AN) fibers through a stochastic model that accounts for stimulus and spike-history effects (Boulet, 2016; Tabibi et al., 2021). The pulses of current represent the electrical signals delivered to cochlear implant electrodes. AN spikes then produce excitatory synaptic conductances that drive voltage activity in the second stage of the model.

The second stage of the model describes an MSO neuron with two separate spatial domains (compartments). The first compartment represents the soma and dendrites, the regions where synaptic inputs are integrated. The second compartment represents the axon initial segment (AIS), the locus of spike-generation. The voltage dynamics of the MSO neuron model are governed by the Hodgkin-Huxley type equations that have been modified for use in auditory brainstem neurons (Rothman and Manis, 2003; Khurana et al., 2011; Goldwyn et al., 2019). The two-compartment structure is adapted from previous models of MSO neurons (Khurana et al., 2011; Goldwyn et al., 2019).

### 2.1 Auditory nerve model

We generated auditory nerve fiber spike trains using a phenomenological, stochastic model of responses to cochlear implant stimuli. The model, developed by Boulet (2016) and also described in Tabibi et al. (2021), randomly generates spikes in response to electrical current pulses that represent the CI stimulus. Our implementation of the model differs from theirs by considering spiking activity on a pulse-by-pulse basis, rather than as a function of time more generally. We detail our approach below. All parameter values, function expressions, and other model features are identical to those used by Boulet (2016).

We describe AN activity level using a state variable *V* (*t*) that represents (phenomenologically) the voltage response of a neuron to stimulation. Voltage is a dynamical variable that responds to input current *I*(*t*) according to the linear differential equation

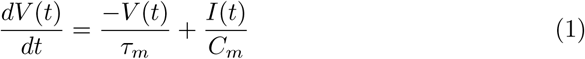

with parameters *τ_m_*= 0.1350 ms and *C_m_* = 0.0714 pF, as in Tabibi et al. (2021). We take the input function *I*(*t*) to be a pulse train of current delivered to a cochlear implant electrode. A single pulse train (to one “ear” of the model) is specified by pulse duration *δ* and interpulse interval Δ, see Fig. 1. Interpulse interval is the reciprocal of pulse rate, and we typically describe a pulse train in terms of the number of pulses per second (pps). For mathematical simplicity, we use monophasic pulses (positive-going current pulse only), whereas pulses in clinical devices are biphasic (positive and negative-going phases of current).

**Fig. 1.**
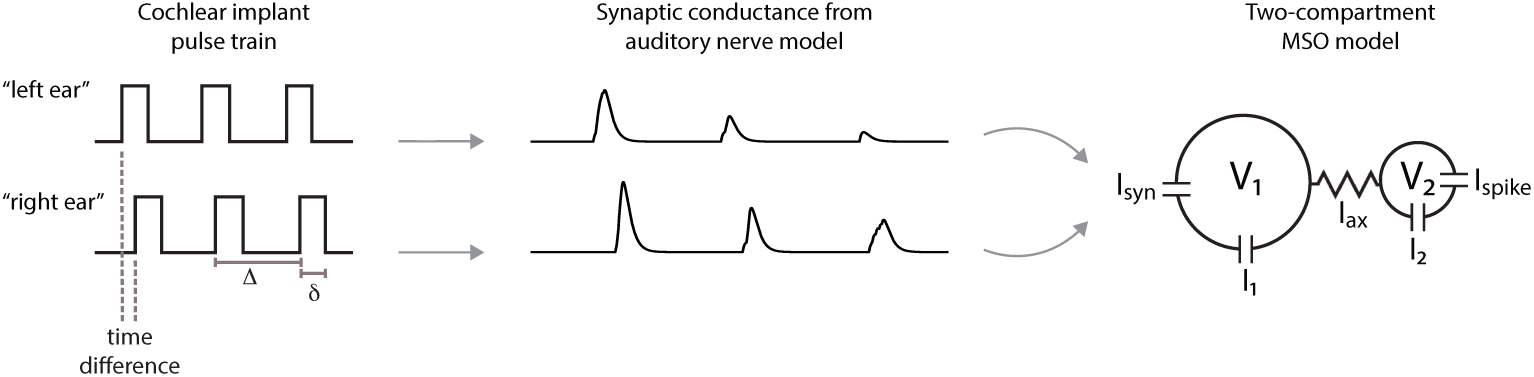
Schematic of model for MSO responses to CI input. Cochlear implant stimuli are represented as pulse of current that drive spiking activity in a model for auditory neurons. Auditory neurons spikes produce excitatory conductance. The combined synaptic conductance from “right ear” and “left ear” inputs enter in the first compartment of the MSO neuron model. Parameters that define the current pulse input are time difference between “right ear” and “left ear” inputs, pulse duration (*δ*), and interpulse interval (Δ, the reciprocal of pulse rate).

The parameter *A_i_* is the amplitude of the *i^th^* pulse. In most simulations we take *A_i_* to be constant across pulses. The exception to this is the final set of simulations in which we vary *A_i_* values according to a sinusoidal function (to represent a signal with an amplitude modulated envelope). In calculations described below that detail the model implementation, we notate the start time of the *i^th^* pulse as *s_i_* and the end time of each pulse as *e_i_*. We refer to *s_i_* as the times of *pulse onsets* and *e_i_* are the times of *pulse offsets*. In simulations of the model, we use two pulse trains of input current that are identical except for a possible time difference between the two signals (Fig. 1). This time difference represents the interaural time difference (ITD) of a sound arriving at the two ears. Our method of quantifying time difference sensitivity in the MSO neuron model is to simulate firing rate activity as a function of time differences between the input pulse trains.

We characterize the dynamics of *V* (*t*) in two distinct time intervals. The first interval is between the start of a pulse at time *s_i_* and the end of that pulse at time *e_i_*. During this interval the pulse is on. The second interval is the time from the end of a pulse (*e_i_*) until the time at the start of the next pulse (*s_i_*_+1_). Pulses are off during this time. The voltage variable increases during the first interval, when the pulse is on with *I*(*t*) = *A_i_*. Voltage decreases during the second interval (the time between pulses), since *I*(*t*) = 0 during these times. We solve the differential equation in Eq. 1 over each of these two intervals using standard methods (Boyce et al., 2021) and obtain the expressions

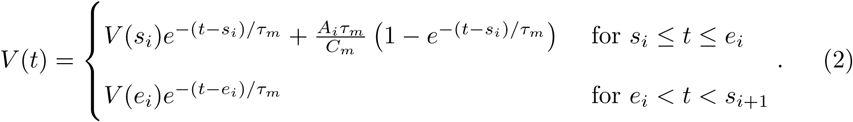

The first equation applies during the pulses and the second expression applies between pulses.

To randomly generate spike times, we transform the value of the voltage variable to a spike probability value. We do this using the firing efficiency function, a concept introduced into cochlear implant modeling by Bruce et al. (1999a). Spike probability as a function of voltage is

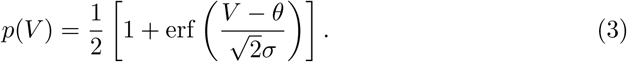

The abbreviation erf is for the error function, the parameter *θ* is the spike threshold (value of *V* at which spike probability is 1/2), and the parameter *σ* determines the variability of firing. Rescaling *σ* by the threshold parameter defines the quantity *ρ* = *σ/θ*. This quantity is known as relative spread (Bruce et al., 1999a) and is frequently used in modeling and neurophysiological studies of AN responses to CI stimulation.

Bruce et al. (1999b) introduced the technique of making threshold *θ* and relative spread *ρ* depend on spike-history in order to describe a refractory period. Boulet, Tabibi, and colleagues expanded on this approach to include dynamics in *θ* and *ρ* that account for observed phenomena of facilitation, accommodation, and spike rate adaptation in auditory nerve fiber responses to CI stimulation (Boulet, 2016; Tabibi et al., 2021). We detail the dynamics of these quantities below, in subsequent sections of Methods. Our implementation of the model differs from Boulet, Tabibi, and colleagues by restricting spikes to occur only at the end of each pulse. To determine spike probability we must only evaluate *V*, *θ*, and *ρ* at the time of pulse offset within each interpulse interval. Our rationale for this choice is that pulse durations are brief (50 *µ*s in our model) and *V* (*t*) rises rapidly during a pulse and decays rapidly after a pulse, so *p*(*V*) is largest at the end of each pulse.

We calculate the voltage at the end of each pulse using Eq. 2. In particular, evaluating the second expression in Eq. 2 at the end-time of a pulse gives

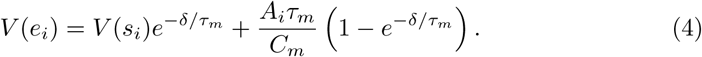

Next, we observe that *V* (*s_i_*) depends on the voltage at the end-time of the previous pulse according to the exponential decay rule *V* (*s_i_*) = *V* (*e_i_*_−1_)*e*^−(Δ−^*^δ^*^)^*^/τm^*. Thus, using the second expression in Eq. 2, we can replace *V* (*s_i_*) with a term that depends on voltage at the ending time of a pulse. From these observations we create an iterative update rule for voltage at pulse offsets:

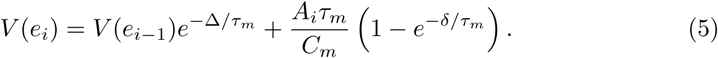

Beginning from an initial value of *V*_0_ = 0, we compute the voltages at the end of each pulse using this formula. We then place these voltage values into Eq. 3 to produce pulse-by-pulse spike probabilities, which we denote *p_i_* (where *i* = 1, 2*, . . .*) labels the pulse number). We reset the voltage variable to zero when spikes occur.

As mentioned above, the values of *θ* and *σ* in the spike probability equation Eq. 3 are also dynamical variables. The values of these paramters change from pulse-to-pulse due to spike history and stimulus history in order to incorporate the effects of refractoriness, facilitation, accommodation, and adaptation. We are calculating spike probabilities at the end of each pulse, thus we require expressions for the values of *θ* and *σ* at the times *e_i_* for *i* = 1, 2, We obtain these expressions by solving relevant linear differential equations, breaking the pulse-to-pulse intervals, in a manner similar to our iterative for *V* (*e_i_*) in Eq. 5. Following Boulet (2016), we use dynamical expressions for relative spread *ρ* and define *σ* through the expression *σ* = *ρθ*.

We define *θ*(*t*) and *ρ*(*t*) as the product of their baseline values (*θ*_0_ = 30 and *ρ*_0_ = 0.04) with dynamic multipliers that implement refractory period, facilitation, accommodation, and adaptation (Boulet, 2016; Tabibi et al., 2021):

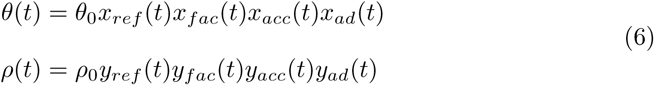

We next describe the dynamics of each of these multiplier terms. The implementation we are presenting here follows the same approach of Boulet, Tabibi, and colleagues and uses their parameter values (Boulet, 2016; Tabibi et al., 2021). The adjustment we are making is to calculated spike probabilities only at pulse offset times.

#### 2.1.1 Refractory period

The refractory period refers to the time immediately after a spike during which a subsequent spike cannot be generated (*absolute* refractory period) or requires additional excitation (*relative* refractory period) (Miller et al., 2001). Let *T* represent the time since the last spike. A second spike cannot be produced for *T* smaller than the absolute refractory period of *t_abs_* = 0.332 ms, meaning spike probability is zero for *T ≤* 0.332 ms and we only use Eq. 3 to calculate spike probabilities when pulse offset time is outside the absolute refractory period. For *T >* 0.332 ms, beyond the absolute refractory period, there is a relative refractory period during which spike probabilities are reduced (larger threshold) and more variable (larger relative spread). In particular, the multiplier terms in Eq. 6 for the refractory effects are

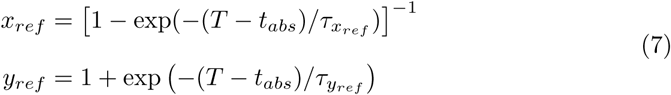

where *τ_xref_* = 0.411 ms and *τ_yref_* = 0.2 ms.

#### 2.1.2 Spike rate adaptation

Spike probabilities also decrease over a longer time scale than the refractory period during periods of high neural activity. This is the phenomenon of spike rate adaptation (Zhang et al., 2007). Our model for adaptation, following Boulet (2016), is to increase the spike rate multiplier variable (and thereby increase the spike threshold) at the time of each spike by an increment of 0.04. In the periods between spikes, this multiplier variable (*x_ad_*) relaxes back to its baseline value of one according to linear dynamics on a slow time scale with time constant *τ_x__ad_* = 50 ms:

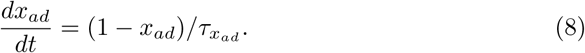

Recall from above that we use *T* to denote the time since the last spike. We also introduce the symbol *X* to be the value of this adaptation multiplier at that previous spike time. We can then solve the above differential equation to find *x_ad_* at any later time (but before the next spike) by initializing the value of *x_ad_* at the time of the spike to be *X* + 0.04 and we find

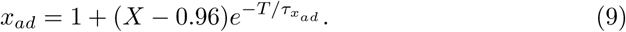

This equation gives the value of *x_ad_* at any time *T* after a previous spike and before the next spike. The same equations govern the multiplier for relative spread, denoted by *y_ad_*.

#### 2.1.3 Facilitation

Facilitation refers to the increase in spike probability that occurs on short time scales if the neuron receives multiple subthreshold inputs that temporally-summate. Facilitation in auditory nerve fiber responses to CI stimulation has been characterized using pairs of pulses. A pulse is delivered at a subthreshold current level to decrease the threshold level of the neuron in response to a second pulse delivered a short time later (Cartee et al., 2000).

The facilitation multiplier is driven by changes in voltage, but with a delay of *δ* (the pulse width) so that the multiplier begins to increase at pulse offset. Contrast this with voltage, which begins to increase at pulse onset. We define the facilitation multiplier *x_fac_* as *x_fac_* = 1 + *z* where the dynamical variable *z* is governed by the linear differential equation

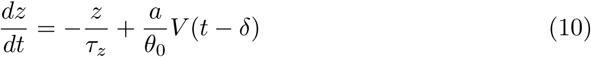

The forcing term for *z* is the delayed value of voltage *V* (*t − δ*) with rescaling of voltage by the parameter *θ*_0_ (the midpoint in the firing efficiency curve) and *a*, the parameter that controls the strength of the facilitation effect. The values for these parameters are *a* = *−*0.15/ms with time constant *τ_z_* = 0.5 ms. The value of *a* is negative in this case because facilitation increases excitability (lowers the threshold value). We solve this in two intervals using the expressions for *V* (*t*) reported in Eq. 2.

Recall, from Eq. 2, that the dynamics of *V* are distinct in two intervals. During a pulse, voltage increases as it is driven by the input current. Between pulses, voltage decays exponentially since it receives no input. To facilitate analysis, we break up the dynamics of *z* into corresponding phases (but shifted, because of the delay term *V* (*t − δ*) in Eq. 10). As introduced previously, use *s_i_* to denote the onset time of the *i^th^* pulse and let *e_i_* = *s_i_* + *δ* be the time at pulse offset. During a short time immediately after pulse offset, that is for the time interval *e_i_ < t < e_i_* + *δ* the value of *z* is driven by the phase of *V* that is increasing in response to the pulse of current. Thus, *z* in a short time interval after the end of a pulse, *z* is governed by the equation

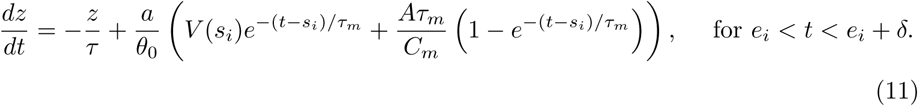

For the remainder of the time until the end of the next pulse, that is for the time interval *e_i_* +*δ < t < e_i_* +Δ, the dynamics of *z* are driven by the exponentially-decaying phase of *V*. Thus, *z* is governed by the equation

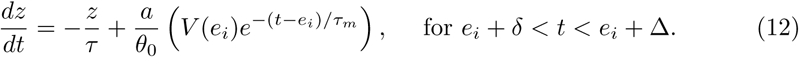

We always choose interpulse intervals to be larger than twice the pulse duration, and so we can first solve Eq. 11 and then use the appropriate value of *z* as the initial value for the next phase of *z* (governed by Eq. 12). In this manner, we can iteratively solve these two differential equations to obtain values of *z* at the end-times of pulses (*e_i_*). Recall, these are the only times at which we allow AN spike generation to occur in our implementation of the model. Our calculation proceeds as follows. First, we use *z*(*e_i_*), the value of *z* at pulse offset, as the initial value for Eq. 11 (with *z* = 0 as the first initial value at the start of the stimulus). We solve this equation at time *e_i_* + *δ* and obtain

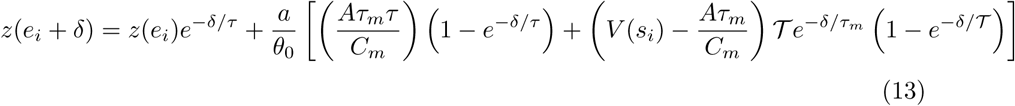

where 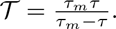

Next, we use this value of *z* as the initial value for Eq. 12. We solve this equation and evaluate *z* at the next pulse offset to find

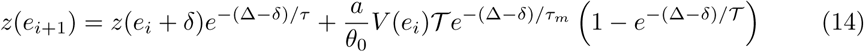

We add one to this value to obtain the value of the threshold multiplier *x_fac_* at the pulse offset, which is the information we require to compute spike probability. The facilitation process resets (*z* = 0) at every spike time and at the end of every pulse (Boulet, 2016). We use the same calculation scheme for the relative spread multiplier *y_fac_*. Parameter values for the relative spread multiplier are *a* = 0.75/ms and *τ* = 0.3 ms. As stated previously, we take all parameter values for the AN model from Tabibi et al. (2021).

#### 2.1.4 Accommodation

On longer time scales than facilitation, multiple subthreshold inputs can have a cumulative effect to gradually increase spike threshold. This is the phenomenon of accommodation (Negm and Bruce, 2014). The multipliers for accommodation are split into two components, quick and slow accommodation. The dynamics of these variables are treated using the same scheme as facilitation. An auxiliary variable (denoted as *z* in the treatment of facilitation, see Eq. 10) has dynamics driven by delayed voltage, and the accommodation multipliers are one plus the auxiliary variable. The accommodation effect makes the AN less excitable (increases spike threshold), thus the scaling parameter *a* is positive for both the quick and slow components of accommodation. For the quick accommodation threshold multiplier *a* = 0.5/ms and *τ* = 1.5 ms, for the slow accommodation threshold multiplier *a* = 0.01/ms and *τ* = 50 ms. Notice that the dynamics of these components are distinguished by their constants, which differ by an order of magnitude. The quick component of the accommodation relative spread multiplier has *a* = 0.75/ms and *τ* = 0.3 ms. There is no slow component for the relative spread multiplier (*a* = 0/ms).

#### 2.1.5 Spike time jitter

As constructed up to this point, our implementation of the AN model restricts spike times to occur only at pulse offsets. While AN spikes are well-timed to the pulses of CI stimulation, they do exhibit a small amount of temporal jitter (Miller et al., 2001). The neural computation we are simulating is coincidence detection between two streams of synaptic inputs, so incorporating this small amount of temporal jitter is an important element of our model. Our approach is to temporally displace each spike time by a random amount. In particular, we add to each spike time a normally-distributed random number with mean zero and standard deviation of 0.1 ms. This standard deviation value is based on physiological measurements of AN spike time jitter in response to CI stimulation (Miller et al., 2001).

### 2.2 Synaptic input model

We use auditory nerve fiber activity as inputs to a biophysically-based computational model of a principal cell in the medial superior olive (MSO). We omit additional, early stages of auditory pathway that are intermediate between the auditory nerve and MSO. These include cochlear nucleus neurons that pass excitatory inputs forward to MSO and medial trapezoid boy neurons that inhibit MSO. Although these nuclei impact MSO tuning (cells in the anteroventral cochlear nucleus can enhance the temporal precision of inputs to MSO neurons (Joris et al., 1994) and inhibitory inputs can adjust the positions of the peaks of ITD tuning curves (Brand et al., 2002; Jercog et al., 2010; Myoga et al., 2014)), our approach captures the essential architecture of the MSO circuit. Specifically, our model describes MSO neurons as neural coincidence detectors of well-timed excitatory inputs originating from both ears.

The synaptic inputs to the MSO neuron model are in the form of excitatory post-synaptic conductances (EPSGs). For each auditory nerve fiber spike, we create an EPSG of the form

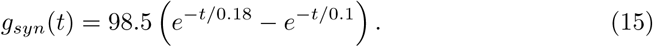

The rise and decay time scales of this double exponential function are taken from reports of *in vivo* MSO recordings (Franken et al., 2015). The amplitude of this unitary EPSG is selected so that a excitatory input depolarizes the soma voltage of the MSO neuron (described below) by roughly 6 mV. We selected this value based on MSO recordings reported in Roberts et al. (2013).

MSO neurons have been described has having few inputs (Couchman et al., 2010), so in our model we use ten independent synaptic inputs for each “side” of the MSO neuron. We vary the time delay of the two input streams (contralateral and ipsilateral, so to speak) to represent sounds that arrive at the two ears with different ITDs. In total, the synaptic input to the MSO neuron is in the form a current

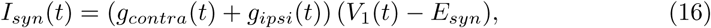

where *g_contra_* and *g_ipsi_* are each composed from the sum of ten synaptic input streams, *V*_1_ is the soma voltage of the MSO neurons (defined next), and *E_syn_* = 0 mV is the reversal potential for synaptic excitation.

### 2.3 MSO neuron model

Principal cells in the medial superior olive are the first nucleus in which bilateral inputs are combined to make fine-timing comparisons to encode sound source locatiion (Grothe et al., 2010, for review). We model these neurons using a two-compartment model that describes separately the dynamics of the soma/dendrite region (first compartment) and the axon initial segment (second compartment). We adapt the model of Khurana et al. (2011), a Hodgkin-Huxley type model with low and high-threshold Potassium currents, spike-generating Sodium current (in the second compartment only), and hyperpolarization-activated cation current; all based on the Rothman-Manis model (Rothman and Manis, 2003). We found the steady state input resistance at the soma of the Khurana et al. model to be 14.8 *M* Ω and the membrane time constant in the soma to be 445 *µs* and retained these values in our model. In earlier work, we discussed how two-compartment models can be parameterized by the relative strengths of forward and backward coupling (Goldwyn et al., 2019). Based on that work, we adjusted parameter values to increase the strength of forward coupling (less voltage attenuation from soma to axon), since stronger forward connection from soma to axon are beneficial for ITD sensitivity (Goldwyn et al., 2019).

We list the parameter values for our model in Table 1 in the column labelled “control.” These parameters result in a forward coupling strength of 0.8 (meaning the passive, steady state voltage in the AIS is 80% of the steady state voltage in the soma for constant current input to the soma) and a backward coupling strength of 0.5 (meaning the passive, steady state voltage in the soma is 50% of the steady state voltage in the AIS for constant current input to the AIS). See Goldwyn et al. (2019) for details and definition of coupling coefficients in two-compartment models. We also adjusted leak reversal potentials in both compartments to achieve a resting potential of -58 mV.

**Table 1.** Parameters for MSO neuron model.

| parameter | control | deprived |
| --- | --- | --- |
| compartment 1 |  |  |
| $c_1$ | 30.1 pF | 30.1 pF |
| $E_{lk,1}$ | -57.9 mV | -57.9 mV |
| $g_{lk,1}$ | 16.5 nS | 16.5 nS |
| $g_{h,1}$ | 76.9 nS | 38.4 nS |
| $g_{KLT,1}$ | 208.6 nS | 104.3 nS |
| $g_{KHT,1}$ | 0 nS | 0 nS |
| $g_{Na,1}$ | 0 nS | 0 nS |
| $g_{M,1}$ | 0 nS | 0 nS |
| compartment 2 |  |  |
| $c_2$ | 14.5 pF | 21.7 pF |
| $E_{lk,2}$ | -59.0 mV | -59.0 mV |
| $g_{lk,2}$ | 3.95 nS | 5.93 nS |
| $g_{h,2}$ | 19.2 nS | 19.2 nS |
| $g_{KLT,2}$ | 52.2 nS | 0 nS |
| $g_{KHT,2}$ | 150 nS | 225 nS |
| $g_{Na,2}$ | 2000 nS | 3000 nS |
| $g_{M,2}$ | 0 nS | 21.9 nS |

The equations that govern the voltage dynamics in the soma/dendrite region (*V*_1_) and the AIS region (*V*_2_) are

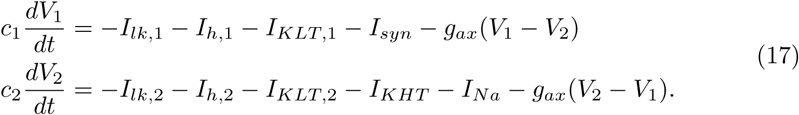

As in Khurana et al. (2011), the expressions for the voltage-dependent currents are *I_lk,i_* = *g_lk,i_*(*V_i_ − E_lk,i_*) for leak currents (we use *i* = 1 or *i* = 2 to label the compartment number), *I_h,i_* = *g_h,i_*(0.65*r_r,i_* + 0.35*r_s,i_*)(*V_i_ − E_h_*) for hyperpolarization-activated cation current, 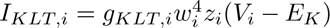 for low-threshold Potassium, *I_KHT_* = *g_KHT_* (0.85*n*^2^ + 0.15*p*)(*V*_2_ *− E_K_*) for high-threshold Potassium current, and *I_Na_* = *g_Na_m*^3^*h*(*V*_2_ *−E_Na_*) for Sodium current. Notice that Sodium and high-threshold Potassium currents are only present in the spike-initiation zone (compartment two). We described the synaptic current term *I_syn_* previously in Eqs. 15 and 16.

All gating variables are governed by differential equations of the form

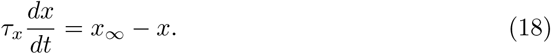

The specific forms of the time constants and equilibrium values are given in Khurana et al. (2011), and we also include these in the Appendix.

### 2.4 Changes to MSO neuron model to describe AIS plasticity

Changes to MSO neurons due to auditory deprivation have not been reported, to our knowledge. We therefore draw on observations from the avian sound localization system to make changes to the MSO model that could represent possible effects of hearing loss on MSO structure and physiology. We refer to the model with parameters altered to reflect the effects of auditory deprivation as the *deprived model*. We contrast the deprived model with the *control model*.

The axon initial segment in neurons in the *nucleus magnocellularis* (NM) of chicks enlarges during periods of auditory deprivation. Specifically, the length of the axon region with dense sodium channels expands, although the distance from the soma to the AIS does not change (Kuba et al., 2010, 2014). Additional studies of AIS plasticity for NM neurons under conditions of auditory deprivation showed that, while there were no changes in the density of Na channels, there were changes to the distribution and make-up of Potassium channels in the soma and AIS (Kuba et al., 2010, 2015). In particular, low threshold Potassium current (Kv1.1 subunit type) diminishes in the soma and AIS. There is a complementary increase in M-type Potassium current (Kv7.2 subunit type) in the AIS during auditory deprivation, indicating a possible reconfiguration of Potassium channels from KLT to M-type. Kuba and colleagues argue that these changes comprise a homeostatic adjustment that maintains neural activity when excitatory synaptic inputs are lost during auditory deprivation (Kuba et al., 2010; Kuba, 2012).

To model possible features of AIS plasticity induced by auditory deprivation, we changed several parameter values. To describe the elongation of the AIS during auditory deprivation, we increased passive parameters (capacitance, leak conductance) in the second compartment by 50%, consistent with an increase in the surface area of that compartment. Sodium channel density has not been observed to change due to auditory deprivation, so we also increased Sodium conductance by 50% to retain the same Na channel density for the larger-sized AIS compartment. To increase excitability – the homeostatic effect observed in experiments – we decreased low-threshold Potassium conductance in the soma by 50%. We obtained similar time difference tuning curves, but with decreased excitability (non-homeostatic) using larger values of Potassium conductance (simulations not shown). We matched the 50% decrease in *g_KLT_* with a 50% decrease in the hyperpolarization-activated cation conductance (*g_h_*) to preserve the balance of these conductances in the soma, as has been observed *in vitro* (Khurana et al., 2011). Lastly, and importantly, based on the experimental findings and modeling work in Kuba et al. (2015), we reduced Potassium conductance in the AIS by 50% and converted the low-threshold Potassium conductance to M-type Potassium conductance.

We list the resulting parameter set for the auditory deprived model in the right column of Table 1. Parameter values that are the same for both compartments and for both control and deprived models are reversal potentials (*E_K_* = *−*106 mV, *E_h_* = *−*47 mV, and *E_Na_* = 55 mV) and the axial conductance connecting the two compartments (*g_ax_* = 56.4 nS).

### 2.5 Numerical methods

We used python code to perform the numerical calculations described above to simulate auditory nerve activity. We used a forward Euler method with time step of 0.002 ms written in C code to solve the differential equations that describe the dynamics of the MSO neuron model. Due to the stochastic nature of the model, most results are reported as mean and standard error of 100 repeated simulations of the mode. To calculate firing rates, we counted spikes over the duration of a 300 ms long simulation, with spikes defined as upward crossings of -20 mV in the *V*_2_ variable (AIS voltage). Code is freely available for inspection and use at https://github.com/jhgoldwyn/auditoryDeprivedMSO.

## 3 Results

### 3.1 AN responses to CI stimulation at varying pulse rates

We used the first stage of the model to create auditory nerve fibers (AN) spike trains in response of to pulses of current that represent cochlear implant stimulation. In the model, AN spiking aligns closely with the ending times of pulses, as shown in the raster plots in the first row of Fig. 2. As we described in Methods, to create temporal dispersion of spike times around the time of pulse offset, we added a small amount of random jitter to each spike time (standard deviation of 0.1 ms). This amount of jitter is not substantial relative to the interpulse intervals, particularly at lower pulse rates (apparent by the sharp peaks in the post-stimulus time histograms (PSTHs) in the second row of Fig. 2). At the lowest pulse rate shown (250 pps, panel A), the interpulse interval is 4 ms. This is sufficiently long to exceed the refractory period. The decrease in firing rate visible over the first 5 ms to 10 ms in this case, and also in response to higher pulse rates (visible in the decreasing amplitudes of peak heights in the PSTHs), is due to the phenomena of adaptation and accommodation. At the highest pulse rate we show (1000 pps, panel C), AN spike times can appear more disorganized across trials than the tightly phase locked responses to low pulse rates. This is particularly the case in the later portions of the responses where adaptation and accommodation take effect.

**Fig. 2.**
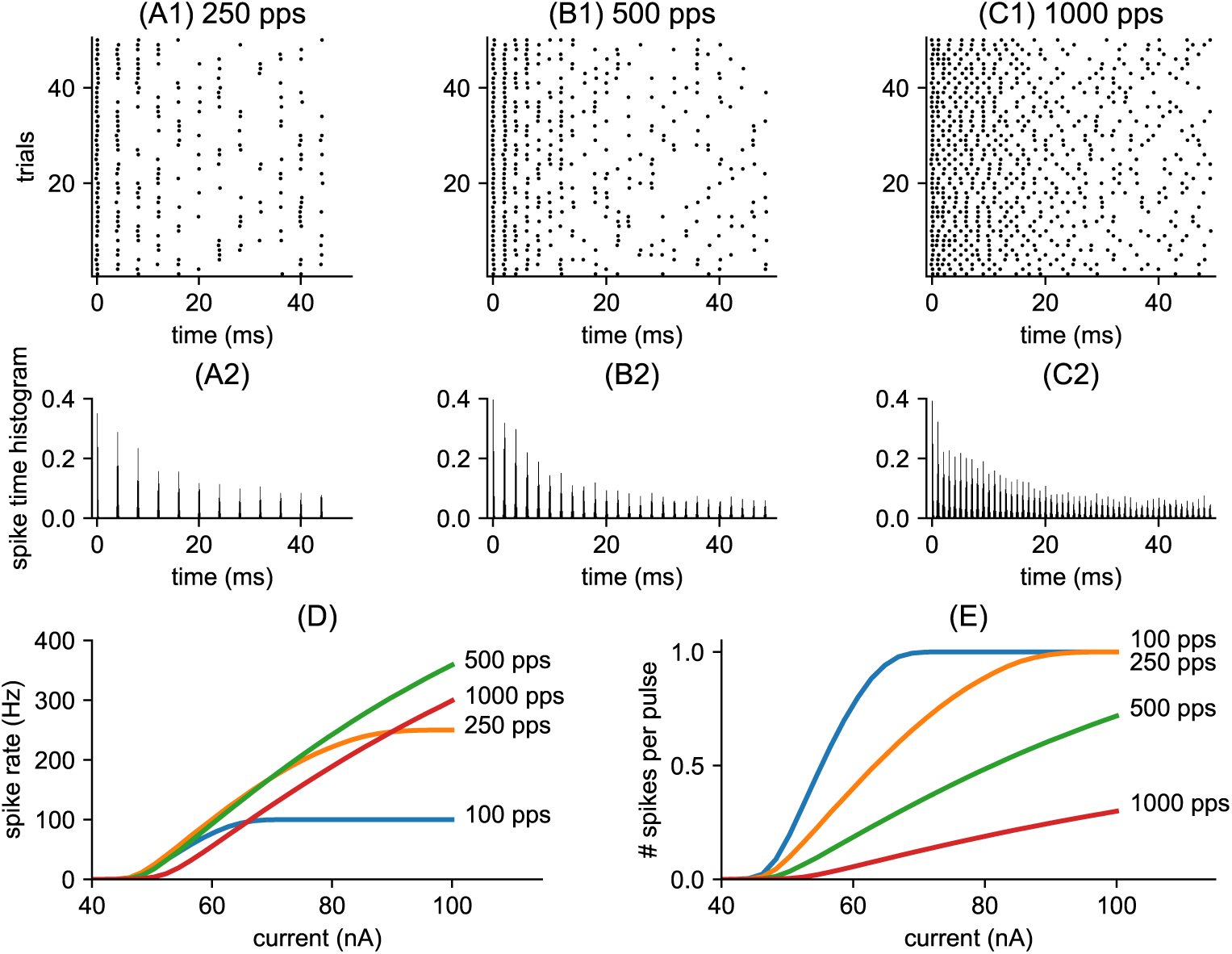
Spiking activity in the nerve (AN) model. Responses to constant amplitude pulse trains. Raster plots show spike times in 50 repeated simulations of the AN model in response to constant amplitude pulse trains **(A1)** pulse rate is 250 pps and pulse amplitude is 55 nA, **(B1)** pulse rate is 500 pps and pulse amplitude is 60 nA, and **(C1)** pulse rate is 1000 pps and pulse amplitude is 75 nA. As specified in the model, spike times are aligned to the pulse offset with a small amount of spike time jitter, as evident in the post-stimulus histograms (PSTHs) shown in the second row for each stimulus setting (PSTHs are computed from 1000 repeated trials using a bin width of 0.1 ms). In **(D):** average spike rate in response to a 300 ms constant amplitude pulse train, plotted as a function of pulse amplitude for various pulse rates. Values are averages from 1000 repeated simulations. AN firing rates saturate at one spike per pulse in responses to lower pulse rate stimuli (100 pps and 250 pps). In **(E)**: the same firing rates as (D) but plotted as firing rate divided by the number of pulses to yield an estimate of average number of spikes per pulse. Spike probability saturates at one spike per pulse in the responses to the lower pulse rate stimuli and remains lower in the responses to the higher pulse rate stimuli.

An important distinction between responses to low-rate and high-rate inputs is the way in which AN firing rate increases with current level (Fig. 2D). In response to low pulse rates, there is a small range of currents over which the spike rate of the AN goes from zero (no spiking) to maximal firing rate (saturating at one spike per pulse). This range is determined by the *σ* in Eq. 3, the steepness of the firing efficiency curve. Thus, at high current levels and lower pulse rates, the AN can fire reliably in response to every pulse. In response to high pulse rate stimuli, firing rates increase more gradually over a larger range of current values. One way to view this distinction is to notice that, in response to high pulse rate stimuli, the AN can achieve high firing rates at relatively lower average number of spikes per pulse. (Fig. 2E).

### 3.2 MSO responses to CI stimulation at varying pulse rates

In the second stage of the model, we use simulated AN spike trains as inputs to a two-compartment model of an MSO neuron. Excitatory synaptic conductances drive voltage responses in the input compartment (*V*_1_ in Eq. 17). This variable represents voltage in the soma (recall we omit dendritic processing from our model). Traces of *V*_1_ responses to constant amplitude pulse trains of various pulse rates are shown in Fig. 3 (thick blue curves). The second compartment of the model represents the site of spike initiation in the axon initial segment. Voltages in this compartment are *V*_2_ in Eq. 17 and portrayed in Fig. 3 as thinner, gray curves. MSO spikes appear as large amplitude events in *V*_2_ traces. We constructed the model to have strong forward coupling and weaker backward coupling (Goldwyn et al., 2019). Consequently, spikes back-propagate weakly into the first compartment. Spike amplitudes are smaller and less visible in *V*_1_ than in *V*_2_.

**Fig. 3.**
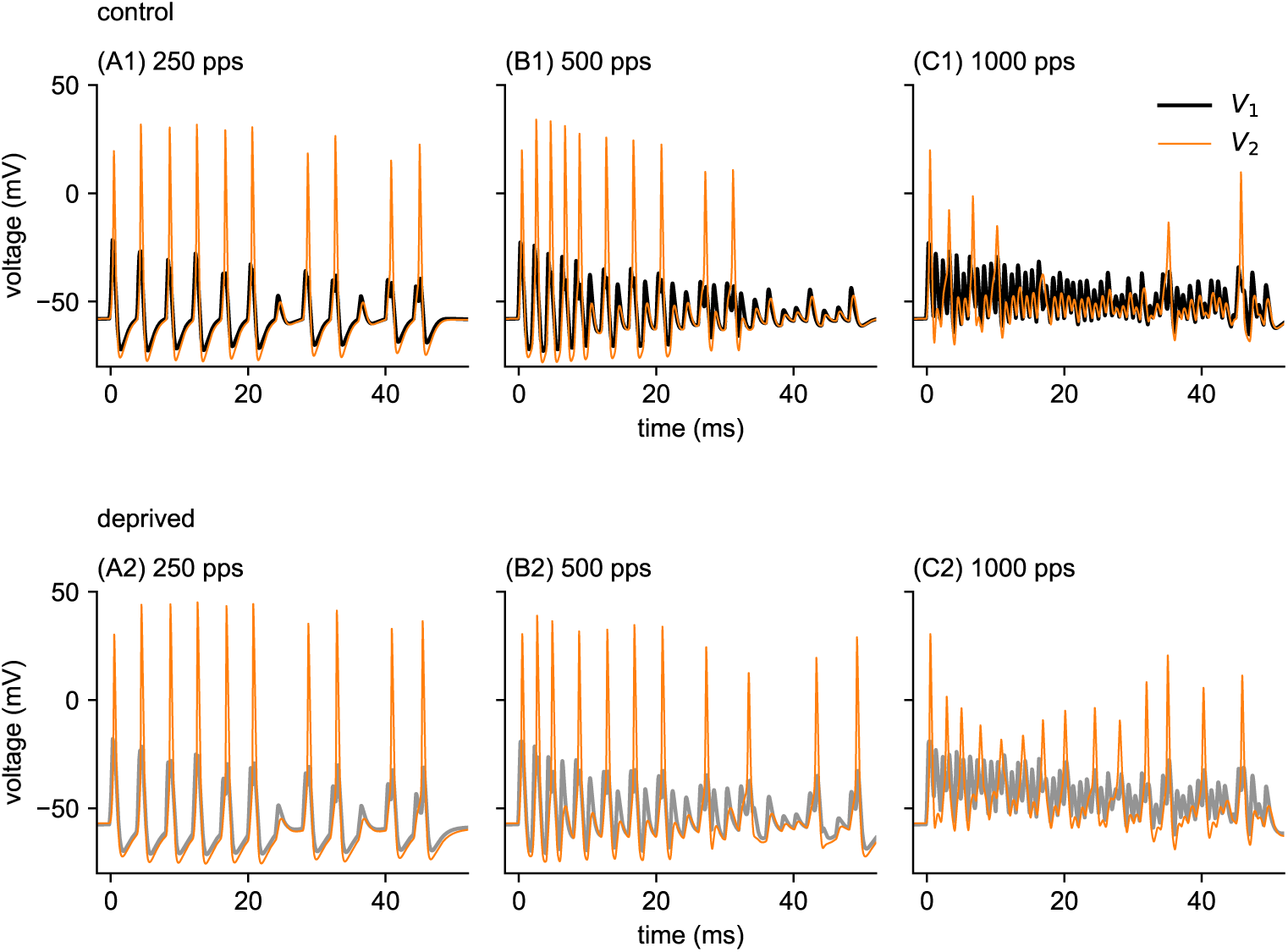
Voltage traces in each compartment of the MSO neuron model in response to cochlear implant stimulation with 0 ms time difference for three pulse rates and amplitudes: Response to the control model are shown in the top row, and responses to the auditory deprived model are shown in the bottom row. **(A)** 250 pps at 54 nA, **(B)** 500 pps at 59 nA, and **(C)** 1000 pps at 81 nA. These stimulus settings were selected so the firing rate is roughly 150 sp/s in the control model in all cases. Thicker black or gray curves are voltages in the soma/dendrite compartment (*V*_1_ in Eq. 17). Thinner orange curves are voltages in the axon initial segment compartment (*V*_2_ in Eq. 17). Spikes are defined as upward crossings by *V*_2_ of the threshold *−*20 mV. Spiking activity entrains to the pulse rate at the low pulse rates (250 pps, left column), but does not spike reliably on a pulse-by-pulse basis at high pulse rates (1000 pps, right column).

For the pulse amplitude used in these simulations, the MSO neuron model entrains reliably to low pulse rate stimuli in both control (Fig. 3A1) and auditory deprived (Fig. 3A2) parameter settings. Spikes occur at the time of (nearly) each pulse in the response to the low pulse rate stimulus (250 pps, Fig. 3A). For higher pulse rates (500 pps in B, and 1000 pps in C), the model cannot generate spikes rapidly enough to entrain to the input on a pulse-by-pulse basis. The firing rate of the control model decreases in response to the highest pulse rate input, so we used a substantially larger-amplitude input current for the high-rate pulse train in these simulations (pulse amplitudes were 54 nA and 59 nA for the 250 pps and 500 pps stimuli, respectively; but 81 nA for the 1000 pps stimulus.

In Fig. 4, we show firing rate responses of the control model (black curves) and the auditory deprived model (gray curves) to constant amplitude, 0 ms time difference pulse trains over a range of current levels and pulse rates. Firing rate responses to the lower pulse rate inputs saturate at the pulse rate (one spike per pulse) at high current levels (Fig. 4A and B). There is no such saturation effect at higher pulse rates for the range of current levels we tested. The MSO neuron control model is most excitable at intermediate pulse rates (500 pps input in Fig. 4) and higher pulse rate stimuli evoke lower firing rates over the same range of pulse amplitudes (Fig. 4D and E).

**Fig. 4.**
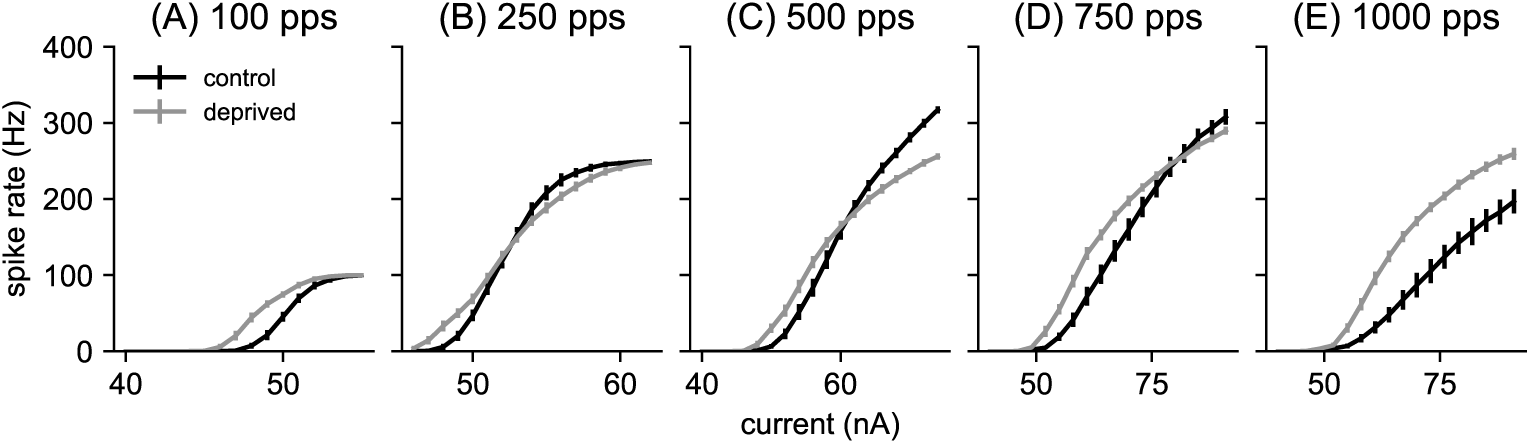
Spike rate for different current strengths, using 300 ms inputs with 0 ms time difference and constant pulse amplitude (current values given on horizontal axis). Pulse rates in each panel are **(A)** 100 pps, **(B)** 250 pps, **(C)** 500 pps, **(D)** 750 pps, and **(E)** 1000 pps. Firing rates increase with current and saturate at the pulse rate for the lower pulse rates (panels A and B). The auditory deprived model is more excitable than the control model (firing begins at lower current levels). Firing rates in the deprived model exceed that of the control model at the lowest and highest pulse rates and at low current levels, but the firing rate of the control model exceeds the firing rate of the auditory deprived model at intermediate pulse rates for large input current levels.

The parameter changes we made to the MSO neuron model (Table 1) reflect changes to axon initial segment observed in auditory brainstem neurons during periods of auditory deprivation (Kuba et al., 2010; Kuba, 2012; Kuba et al., 2014, 2015). These changes have been interpreted as a homeostatic response that compensates for the loss of synaptic drive with an increase in intrinsic excitability (Yamada and Kuba, 2016). Consistent with this notion, we find our auditory deprived model to be more excitable than the control model over a range of pulse rates (gray curves in Fig. 4). This is true in two senses. First, the current level at which spiking activity initaties is lower in the auditory deprived model than the control model across all pulse rates. Second, at all current levels for the lowest pulse rate (Fig. 4A) and the highest pulse rate stimuli (Fig. 4E), and at lower current levels for the intermediate pulse rates (Fig. 4B-D), the auditory deprived model fires at a higher rate than the control model. In response to the highest current values and intermediate pules rates there appears to be some firing rate saturation in the deprived model that allows the firing rate of the control model to exceed the firing rate of the deprived model (Fig. 4B-D, at high current levels).

### 3.3 Time-difference tuning is degraded in the auditory deprived model

To assess time-difference sensitivity in our MSO neuron model, we simulate responses to inputs with time delays between the pulse train inputs to the AN model. This time-shift between the two pulse trains represents an interaural time difference between two sounds arriving at the two ears of a listener using bilateral implants. MSO neurons are coincidence detectors, known to fire maximally to inputs that are aligned in time (Svirskis et al., 2004; van der Heijden et al., 2013). The extent to which firing rates decrease as the time differences between to the two inputs increases is a straightforward measure of the time difference sensitivity of the model neuron.

We find time difference sensitivity, indicated by firing rates that depend on time difference, in the responses of the control model to pulse rates below 1000 pps (black curves in Fig. 5). The decrease in firing rate with increasing time difference is most pronounced at the intermediate pulse rate (500 pps, panel C in the figure).

**Fig. 5.**
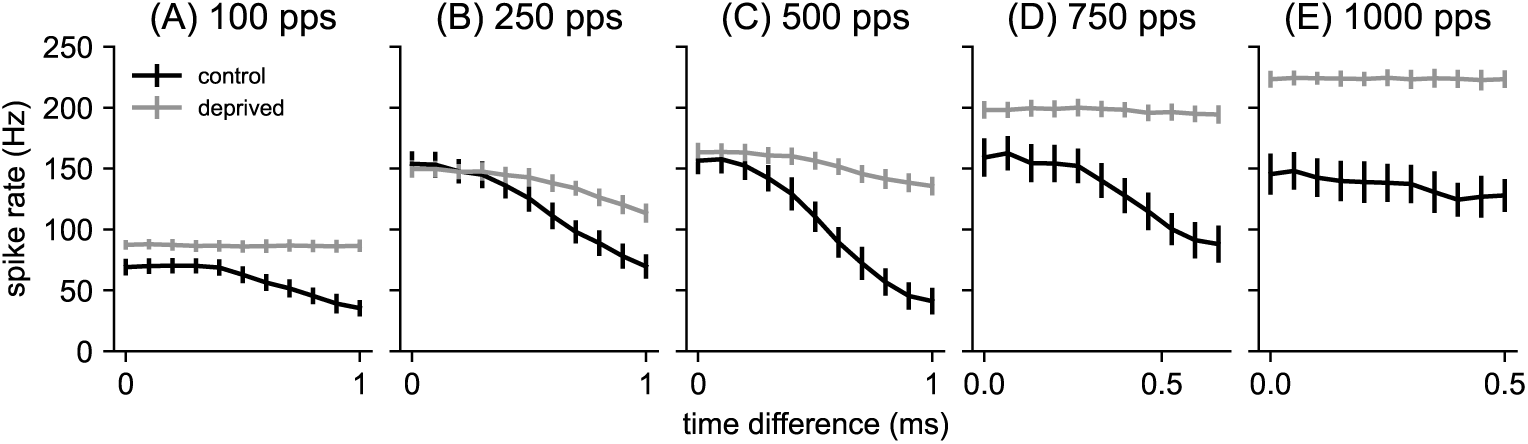
Time difference tuning curves for control model and auditory deprived model for varying pulse rates: **(A)** 100 pps at *A* = 51 *nA*, **(B)** 250 pps at *A* = 53 *nA*, **(C)** 500 pps at *A* = 60 *nA*, **(D)** 750 pps at *A* = 70 *nA*, **(E)** 1000 pps at *A* = 80 *nA*. Spike rate values on the vertical axis are mean of 100 repeated simulations and error bars are standard deviation of those values. Time difference on the horizontal axis varies from 0 (in phase) to 1 ms, except in the case of higher pulse rates (500 pps, 1000 pps). In these cases, the largest time difference possible is half the interpulse interval. Spike rates decrease more as time difference increases for the control model than the auditory deprived model, indicating greater time difference sensitivity for the control model.

Overall, firing rates in the auditory deprived model are flatter with respect to changes in time difference, indicating that the parameter changes we made to create the auditory deprived model degraded time difference sensitivity. Indeed, firing rate responses of the deprived model to the lowest pulse rate (100 pps, Fig. 5A) and to the two higher pulse rates (750 pps in Fig. 5D and 1000 pps in Fig. 5E) are constant with respect to changes in the relative timing of the two pulse train inputs. These tuning curves were computed at specific input levels (see caption of Fig. 5). Next, we show that the same result holds over a large range of current levels.

In Fig. 6 we display time difference sensitivity across a large range of stimulus conditions by plotting firing rates at 0 ms time difference with respect to current level (same curves as in Fig. 4), along with firing rate responses to stimuli with larger time difference. We used 1 ms for this larger time difference in cases for which this did not exceed half the interpulse interval. For the pulse rates that exceeded this limit, we set the maximum possible time difference to be half the interpulse interval (Fig. 4D, E). We plotted firing rates in response to pulse trains with these larger time differences on the same axes, and shaded the region in between to display the range of firing rates produced by the model neuron between these two extreme time difference values.

**Fig. 6.**
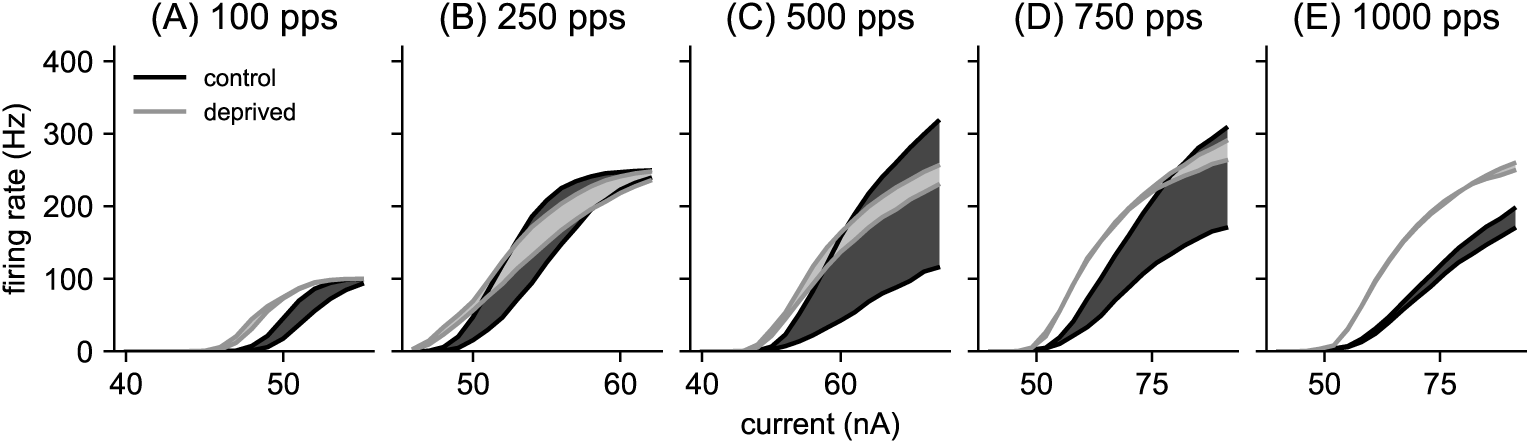
Summary of firing rate sensitivity to time differences. Spike rates of MSO neuron model to constant amplitude pulse trains for various pulse rates: **(A)** 100 pps pulse rate, **(B)** 250 pps pulse rate, **(C)** 500 pps pulse rate, **(D)** 750 pps pulse rate, and **(E)** 1000 pps pulse rate. Maximal firing rates occur for inputs with 0 ms time difference (upper solid curves). Minimal firing rates occur for larger time differences (lower solid curves, using 1 ms time difference for lower pulse rates and half the interpulse intervals for 750 pps and 1000 pps). Values are averages over 100 repeated trials. The shaded regions show the range of possible firing rates between these upper and lower firing rates. Time difference sensitivity, characterized here as the range of firing rates (vertical height of shaded region), is greatest at intermediate pulse rates and greater for the control model than the auditory deprived model.

The control model (dark gray regions) exhibits a small amount of time difference sensitivity in its responses to low pulse rate stimuli (100 pps in Fig. 6A) and high pulse rate stimuli (1000 pps in Fig. 6E). Responses to the intermediate pulse rate, however, show large differences between inputs with no time difference and inputs with a large time difference (1 ms, in this case). The response regions of the auditory deprived model (light gray regions) are substantially narrower than the response regions of the control model (dark gray regions) across all pulse rates. Thus, by this measure of firing rate differences, the auditory deprived model is less sensitivity to time differences than the control model.

### 3.4 Envelope time-difference sensitivity in resposnes to high-rate pulse trains with time-varying pulse amplitude

In typical listening conditions for CI users, the amplitude of each pulse changes over time due to the dynamically changing intensity of sounds. It is of interest, then, to also study time difference sensitivity in the case of pulse amplitudes varying with time. To this end, we simulated responses to high-rate pulse trains with sinusoidally-varying pulse amplitudes. We exhibit time difference tuning as above, shading the firing rate regions between firing rates to inputs with and without a time shift. We used a high-rate pulse train that, if unmodulated, evokes responses in that are constant with time difference (Fig. 7A, note the maximum time difference in this case is 0.16 ms, half the interpulse interval).

**Fig. 7.**
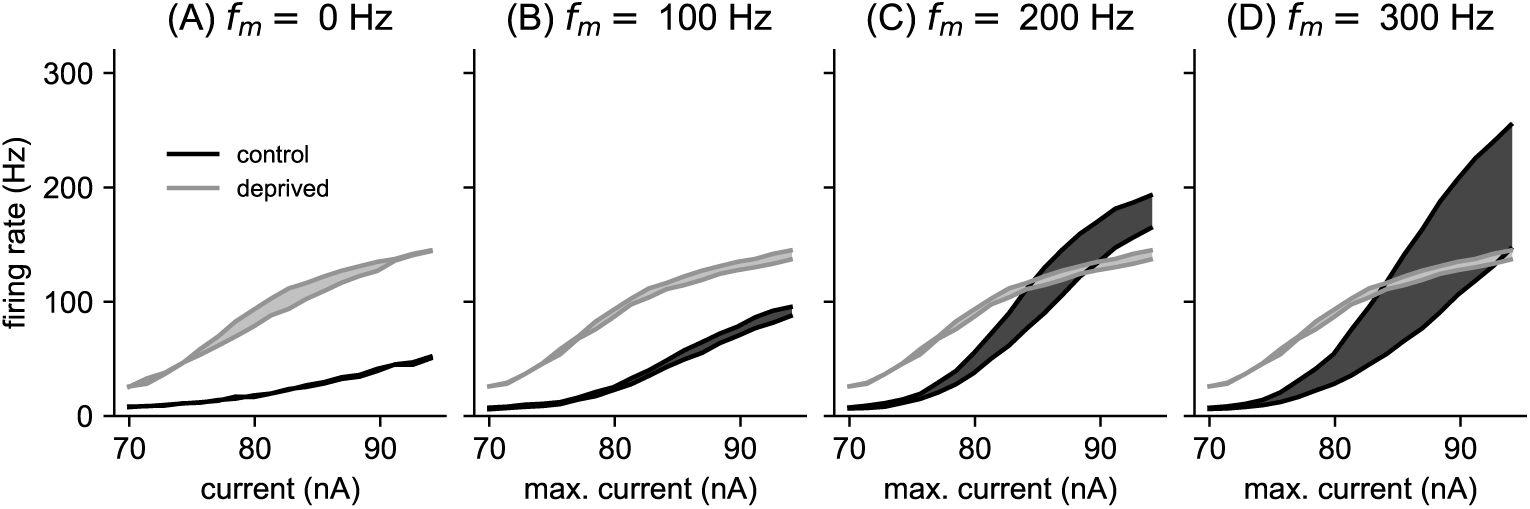
Envelope time-difference sensitivity. Spike rates of MSO neuron model to 3000 pps pulse trains with sinusoidally amplitude-modulated pulse amplitudes. The sinusoidal oscillation of pulse amplitudes varies between a minimum level of 70 nA and a maximum level shown on the *x*-axis. The modulation frequencies are: **(A)** 0 Hz, unmodulated pulse train, **(B)** 100 Hz modulation frequency, **(C)** 200 Hz modulation frequency, and **(D)** 300 Hz modulation frequency. Maximal firing rates occur for inputs with 0 ms time difference (upper solid curves). Minimal firing rates occur for non-zero time difference (lower solid curves, 0.16 ms for the unmodulate pulse train in (A), and 1 ms time difference in all other cases). Values are averages over 100 repeated trials. The shaded regions show the range of possible firing rates between these upper and lower firing rates. Time difference sensitivity, characterized here as the range of firing rates (vertical height of shaded region) is greater for the control model than the auditory deprived model for the higher modulation frequencies (C, D).

If this high-rate pulse train is the carrier signal for a pulse train with slowly-varying pulse amplitudes, however, the model does exhibit time difference sensitivity (Fig. 7B-D). We used 1 ms as the maximum time difference in the case of the amplitude-modulated pulse trains. We find that the control model exhibits time difference tuning, with increasing time difference sensitivity (larger shaded responses regions) at the higher modulation frequencies that we tested. The deprived model, in contrast, does not exhibit similar levels of time difference sensitivity to amplitude-modulated pulse trains. Even at the highest modulation frequency we tested (300 Hz) the auditory deprived model does not exhibit substantial firing rate differences between inputs with 0 ms time difference and inputs with larger time differences (Fig. 7D).

### 3.5 Firing type differences in responses to constant current inputs

We demonstrated that time difference sensitivity in the deprived model is degraded relative to the control model. By this we mean that time difference sensitivity, as measured by the difference between maximum and minimum firing rates (the vertical height of the shaded regions in Fig. 6), is greater for the control model across a large range of input current values. A possible question regarding this result is whether it is simply due to over-excitability in the deprived model. In other words, perhaps the deprived model fires at high rates, generally, including in response to inputs with non-zero time difference. While the deprived model is more excitable, in line with the physiological reports on which we based our model, this alone does not explain why time difference sensitivity is diminished in the deprived model. In particular, if we were to compare tuning curves at fixed firing rates, rather than comparisons at fixed current levels (as in Fig. 5), we would still see greater time difference sensitivity in the control model. This can be deduced by comparing the widths of the response regions in Fig. 6 at locations at which the firing rates are matched, even if the current level is not.

What, then, underlies the degraded time difference sensitivity in the auditory deprived model? We answer this question by pointing out that the type of spiking dynamics changes due to AIS plasticity, in addition to the degree of excitability or average firing rate. By this we mean that control model is phasic firing – it responds to a constant-level current injection with (at most) a single spike at the stimulus onset (Fig. 8A). In contrast, the deprived model fires repetitively to constant-level current inputs (Fig. 8B). This type of response is known as tonic firing.

**Fig. 8.**
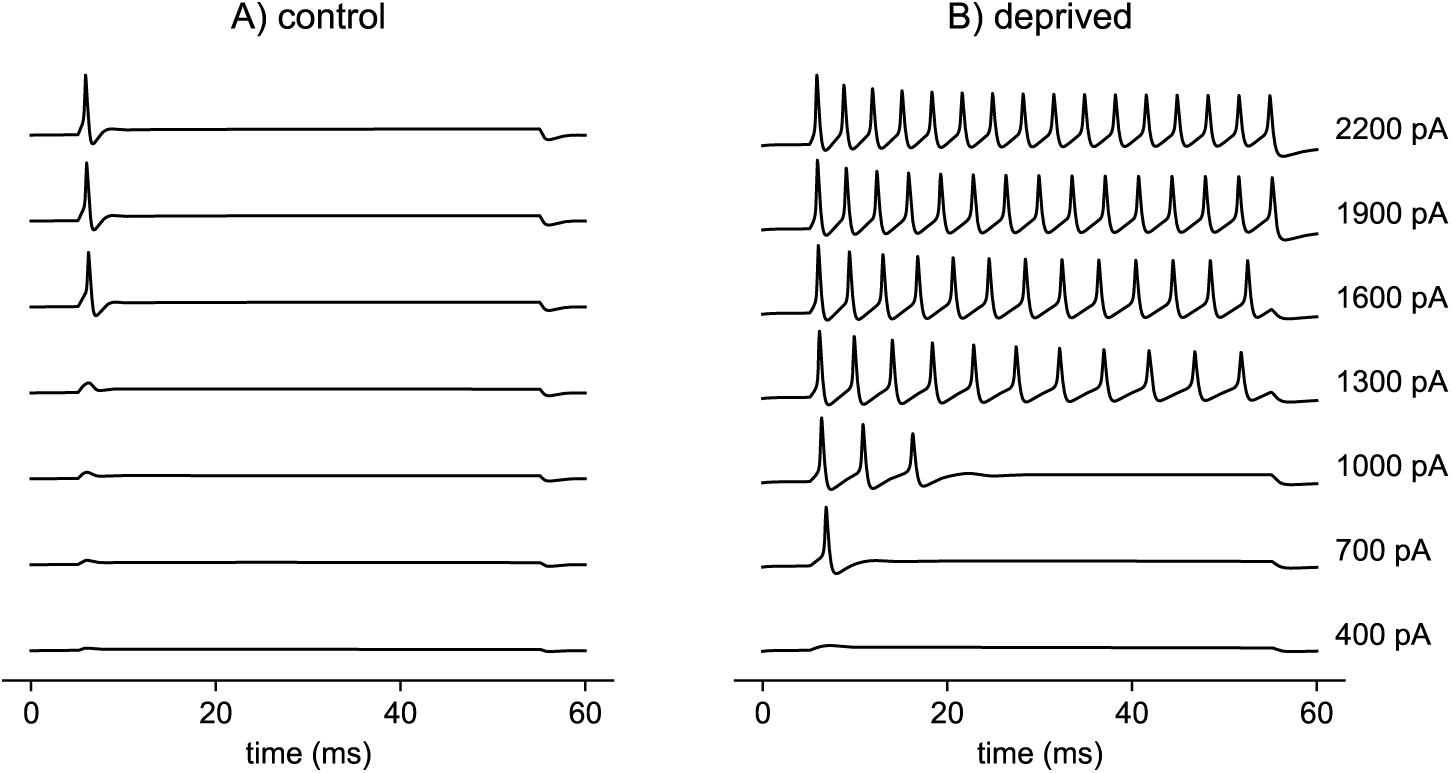
Firing type is phasic for control model and tonic for auditory deprived model. AIS voltage response (*V*_2_, voltage in the second compartment) in response to steps of current injections (50 ms duration), with current level ranging from 400 pA to 2200 pA in steps of 300 pA. In **A** (left column): control model responds with a single spike at stimulus onset (phasic firing type) to inputs above the threshold level of approximately 1360 pA. In **B** (right column): deprived model responds with repetitive firing (tonic firing type) to inputs above approximately 1250 pA. The threshold for a single onset spike in the auditory deprived model is 500 pA (not shown).

MSO neurons are phasic firing cells (Svirskis et al., 2002). Phasic firing is commonly associated with temporally-precise coincidence detection (Meng et al., 2012; Huguet et al., 2017) and therefore would be beneficial for sound localization. By changing the firing type of the model from phasic to tonic, the model loses this dynamical specialization for temporal precision. As a result, time difference sensitivity in the control model exceeds that of the auditory deprived model, as we have shown throughout. The parameter manipulations that caused this change in firing type include reductions to the low-threshold Potassium and *h*-type conductances in the soma and transformation of low-threshold Potassium conductance in the AIS compartment to M-type Potassium (see Table 1). We made these changes based on the experimental and modeling study of AIS plasticity in NM neurons under conditions of auditory deprivation in Kuba et al. (2015).

## 4 Discussion

The use of cochlear implants by individuals with severe to profound sensorineural hearing loss can increase their speech understanding, particularly in noise-free environments (Krueger et al., 2008; Gaylor et al., 2013). Individuals with hearing loss in both ears who use a second implant (bilateral CI) can achieve higher rates of speech understanding than users of a single implant (Litovsky et al., 2006). Bilateral implants, by providing auditory inputs to both ears, have potential to improve perception of the aspects of sound that require binaural processing. Sound source location is one such percept. Two of the primary cues for sound source location are interaural level differences (ILDs) and interaural time differences (ITDs). While bilateral implant users can experience improved sound source localization relative to users with a single CI (Brown and Balkany, 2007), sound source localization and spatial hearing remain a deficit for CI users relative to normal hearing listeners (Grantham et al., 2007).

When evaluating the limitations of CIs, it is essential to consider the relative extents to which CI users’ hearing outcomes are limited by the auditory periphery and central stages of the auditory system. The factors that affect the effectiveness of the implant at the periphery are numerous. These include, among other factors, the signal processing algorithms that convert acoustic inputs to electrical stimuli, the engineering design of the devices, damage to the cochlea during surgery, and pathologies in the cochlea and auditory nerve depending that can depend on the etiology of hearing loss (what has been termed the electrode-neuron interface (Bierer, 2010)). While these factors certainly affect hearing outcomes for CI users, it is also possible that hearing outcomes are impaired by more central stages of neural processing in auditory pathway.

Hearing development passes through critical periods of development that may be disrupted by hearing loss early in life (Kral, 2013; Persic et al., 2020). Hearing loss later in life and even into adulthood can also lead to neural plasticity that reshapes auditory cortex (Eggermont, 2008). In addition to these wide-ranging, circuit-level changes to the auditory system during hearing loss, periods of auditory deprivation in birds (as a model for hearing loss) can induce cellular-level physiological changes in individual neurons. Kuba et al. (2010) found axon initial segments in the nucleus magnocellularis of chicks grew during periods of auditory deprivation, and Kim et al. (2019) found the AIS in the medial nucleus of the trapezoid body in deaf mice to be larger and more distant from the soma than in normal-hearing mice.

AIS plasticity has not been studied in the MSO of humans, to our knowledge. We used Kuba’s studies of NM and NL (nuclei in the sound source localization circuit of birds), as motivation to model similar changes in MSO neurons. Specifically, we compared a control model of MSO neural dynamics to a modified model, which we called the (auditory) deprived model. Based on the experimental findings and computational modeling by Kuba and colleagues, we constructed the auditory deprived model by reducing the amount of low-threshold Potassium conductance in the soma (Kuba et al., 2015) (first compartment of our two-compartment model), transforming the lowthreshold Kv1.1 Potassium channel into to M-type Kv7.2 Potassium channels in the AIS region (Kuba et al., 2015) (second compartment in our two-compartment model), and increasing the size of the AIS (Kuba et al., 2010).

We consistently found the deprived model to be more excitable than the control model (Figs 4 and 8) with poorer time difference sensitivity (as measured by the range of maximum to minimum firing rates in time difference tuning curves, see Figs 5 and 6). Although neither model was sensitive to small differences in the timing of high-rate, constant-amplitude pulse trains (Fig. 6E), the control model exhibited greater time difference sensitivity to amplitude-modulation envelopes of high pulse rate carriers than the deprived model (Fig. 7).

### 4.1 Relation to previous models

Our approach to simulating MSO response to CI stimuli is similar to a previous modeling study by Chung et al. (2015). That paper builds on earlier work (Colburn et al., 2008) by incorporating a more detailed auditory nerve model. Similar to the model we used (Boulet, 2016; Tabibi et al., 2021), Chung *et al*. model auditory nerve responses to CI stimuli as a stochastic process governed by a voltage-like variable that evolves with linear dynamics and a probabilistic spike generation rule based on a dynamic threshold with additive Gaussian noise. Spike threshold in their model is dynamic to describe a refractory period (see also (Bruce et al., 1999a,b)) and also includes spike rate adaption (Nourski et al., 2006). The model we present follows this same general approach and includes refractory and adaptation effects. In addition, we include two additional effects: accommodation (a long time-scale decrease in spike probability) and facilitation (temporal summation of subthreshold inputs to increase spike probability on a short time-scale). These modifications, adapted from the work of Boulet, Tabibi and colleagues (Boulet, 2016; Tabibi et al., 2021), enhance the model by capturing additional nuances of neural responses to CI stimuli.

Chung and colleagues found that their MSO neuron model produced little to no spiking activity in response to unmodulated pulse trains at rates of 500 pps and higher. They modified their model to produce higher firing rates by strengthening synaptic inputs and increasing the speed of membrane dynamics. No such manipulation was needed in our model to drive MSO activity in response to high-pulse rate stimuli. That being said, time difference sensitivity in our model degrades at higher pulse rates, with no significant time difference sensitivity in responses to 1000 pps stimuli. This limitation at high pulse rates is consistent with a recent experimental study showing ITD sensitivity in adult rats implanted with bilateral CIs for pulse rates as high as 900 pps, but not for 1800 pps pulse trains (Buck et al., 2023). It is possible that by using fewer, stronger synaptic inputs and faster membrane dynamics, we could obtain time difference sensitivity in response to higher-pulse rate stimuli, but we have not tested that in our simulations. In our conversion of the control model to auditory deprived model, we did alter the speed of membrane dynamics, but made no changes to synaptic inputs (Kuba et al., 2010). Following observations by Kuba and colleagues and their modeling study (Kuba et al., 2015), we removed a portion of the low-threshold Potassium current (Kv1.1-type channels) from the soma to create the auditory deprived model. This has the effect of increasing the input resistance at the soma and slowing the soma dynamics. Our, therefore, results show the typical result that faster membrane dynamics improve time difference sensitivity and neural coincidence detection (Kempter et al., 1998).

The changes we made to the second compartment, to reflect the processes of AIS plasticity during sound deprivation, enlarged the surface area of this compartment and replaced low-threshold Potassium with M-type Potassium conductance. By enlarging the spike-initiation region in the deprived model, we increased backpropagation of voltage from the AIS into the soma. In a previous work, we described this as an increase in backward coupling (Goldwyn et al., 2019). In that prior study, we found that the greatest time difference sensitivity occurs for with strong forward coupling and weak backward coupling between the soma and axon. The effect of auditory deprivation in our model is to move the soma-axon coupling out of this optimal configuration. Removing the low-threshold Potassium conductance from the spike-generating region, where it functions as a source of dynamic negative feedback (Huguet et al., 2017), also diminishes time difference sensitivity in models of neural coincidence detection (Goldwyn et al., 2019).

### 4.2 Limitations of the model

In the first stage of our model, we used an up-to-date model of temporal response properties of auditory nerve fibers to CI stimulation (Boulet, 2016; Tabibi et al., 2021). The accommodation component of this model distinguishes it from previous models constructed along similar principles (Bruce et al., 1999b; Goldwyn et al., 2012). Accommodation is particularly relevant for modeling high-rate pulse trains as it strongly suppresses spiking under these conditions (Fig. 2). A notable absence in our CI and AN modeling is that we do not describe how the spatial spread of electric fields in the cochlea activates a distributed population of AN fibers, as has been done by others models (Rattay et al., 2001; Cohen, 2009; Goldwyn et al., 2010; Kalkman et al., 2015; Malherbe et al., 2015, for examples). Understanding how population-level auditory nerve activity flows into the MSO circuit remains an important question. On this point, some have suggested that MSO neurons may receive inputs from tonotopically misaligned input (Shamma et al., 1989; Bonham and Lewis, 1999; Day and Semple, 2011). For bilateral CI users, there is almost certainly some tonotopic mismatch in the inputs to binaural circuits because the surgical placements of electrode arrays in two different ears are not perfectly aligned. Mismatched place-of-stimulation does impair lateralization of sounds by binaural CI users (Kan et al., 2013). Future modeling studies of MSO responses to CI stimulation could consider these spatial and population-level effects.

In the second stage of our model, we used a two-compartment model of MSO neural dynamics, similar to previous MSO studies (Khurana et al., 2011; Goldwyn et al., 2019). With the two-compartment structure we can separate the integrative region of the cell where inputs arrive (soma and dendrite regions, represented by the first compartment) from the spike-generating region (in the AIS, represented by the second compartment). Our goal was to understand how AIS plasticity affects time difference sensitivity in the MSO. With this model structure, we represented the AIS and modified its parameters in accordance with AIS plasticity in a straightforward way. Morphological features that we did not include in our model also affect ITD tuning in MSO neurons, including their bipolar dendritic structure (Carr et al., 1998; Dasika et al., 2007; Mathews et al., 2010), and the distribution of mutiple spike initiation sites along its axon (Lehnert et al., 2014). If further evidence regarding AIS plasticity in MSO neurons becomes available, our model could be elaborated to incorporate additional morphological detail.

The connection from the auditory nerve stage to the MSO stage is direct in our model. We omitted nuclei intermediate to these two stages. In response to acoustic stimuli, the anteroventral cochlear nucleus (AVCN) can enhance phase-locking so that spike timing of these cochlear nucleus neurons is improved relative to spike timing in the auditory nerve (Joris et al., 1994). While this enhancement may be important for the temporally-precise dynamics that sound localization requires, in the case of cochlear implant stimulation, the degree of phase locking in the auditory nerve is already exceptionally high (Miller et al., 2008). Phase-locking enhancement by the AVCN may be less relevant in the case of cochlear implants.

Lastly, we did not include inhibitory inputs to MSO neurons, instead focusing on the “excitatory-excitatory” coincidence detection function of MSO neurons. Given the great interest in how inhibitory inputs impact sound localization processing in the MSO (Brand et al., 2002; Grothe, 2003; Jercog et al., 2010; Roberts et al., 2013; Myoga et al., 2014), future work could include this pathway. Elements of the inhibitory pathway also exhibits activity-dependent refinement (Kapfer et al., 2002) and AIS plasticity during auditory deprivation (Kim et al., 2019). Future modeling could study how the inhibitory components, and possible plasticity mechanisms therein, impact sound localization by bilateral CI users.

### 4.3 Implication for sound localization by binaural cochlear implant users

Cochlear implants can improve speech understanding for individuals with profound to severe hearing loss, and yet challenges remain. At the periphery of the auditory pathway, there are factors that can limit CI effectiveness. These include the signal processing performed by the devices, the coarse nature of neural activation by CI stimulation (spatially and temporally), and degradation of the electrode-neuron interface (death or degeneration of AN fibers, for instance). In this study, we showed that neural processing in downstream stages of the auditory pathway may also limit the effectiveness of CIs. In particular, we adjusted biophysical and morphological properties of MSO neurons (AIS plasticity) and found substantial effects on the temporally-precise neural computations that underly sound source localization.

A specific insight that we emphasize is that processes that may appear homeostatic in nature may alter the dynamics and function of neural systems in consequential ways. Changes to a neuron’s properties that increase excitability (to compensate for diminished excitatory drive under conditions of auditory deprivation), can transform the firing type of a neuron. Changes that appear homeostatic with respect to firing threshold or average firing rate can alter the functioning of neurons and neural circuits.

Two features of our simulation results have direct bearing on the high-pulse rate stimulation strategies used in contemporary CIs. First, we observed in the model a decrease in time-difference sensitivity at high pulse rates (1000 pps and higher). The range of available ITDs becomes increasingly limited at high pulse rates. For a 1000 pps pulse train, for instance, the maximum possible ITD is 0.5 ms. More generally, the loss of ITD sensitivity at high pulse rates is consistent with the typical understanding of the MSO as a nucleus that extracts ITD information from low-frequency sounds. Second, MSO neurons can be sensitive to time-differences conveyed by high pulse rate stimuli when those time differences are contained in slowly-varying pulse amplitudes (envelope ITDs, as opposed to fine-structure ITDs). This result is consistent with a previous modeling study of envelope ITD sensitivity in MSO responses to CI stimuli (Colburn et al., 2008). It also supports physiological evidence that the MSO neuron, although typically thought of as a low-frequency nucleus, is sensitive to ITDs in the envelopes of high-frequency inputs (Gai et al., 2014).

Lastly, it is important to point out that we have considered the possibility that there is AIS plasticity in MSO neurons during periods of auditory deprivation (envisioned here as the periods of hearing loss that precede implantation), but we have not considered the possibility that AIS plasticity may occur post-implantation. After CIs are turned on and evoke neural activity, can they initiate homeostatic processes that reduce neural excitability and reverse the AIS plasticity processes that takes place during auditory deprivation? Understanding the plasticity that occurs post-implantation, and the extent to which such processes can improve hearing outcomes for CI users, is an intriguing line of future work.

## Ethical approval

Not applicable. No experimental data was used.

## Contributions

Conceptualization (JHG), data curation (JHG), funding (JHG), investigation (AJ, SX, IF, JHG), methodology (AJ, SX, IF, JHG), software (AJ, SX, IF, JHG), supervision (JHG), visualization (AJ, SX, IF, JHG), writing – original draft (AJ, SX, IF, JHG), writing – review and editing (AJ, SX, IF, JHG).

## Data availability

python and c code for model simulation and data for reproducing figures are available at https://github.com/jhgoldwyn/auditoryDeprivedMSO.

## Appendix

We used the following equations for the dynamics of the gating variables in the MSO model. All gating variables evolve according to a differential equation of the form

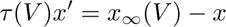

All equations listed below have been used in previous modeling of MSO neurons (Khurana et al., 2011; Goldwyn et al., 2019). Most were first introduced by Rothman and Manis in their study of the cochlear nucleus (Rothman and Manis, 2003). We adjusted kinetics as necessary for a temperature of 35^◦^*C*. We include an M-type Potassium channel in our model because studies of AIS plasticity have shown up-regulation of the Kv7.2 channel type (Kuba et al., 2015). We use the model for M-type Potassium given in that same reference (Kuba et al., 2015).

### Low-threshold Potassium current

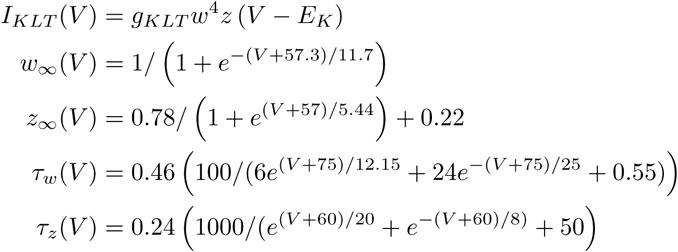

### High-threshold Potassium current

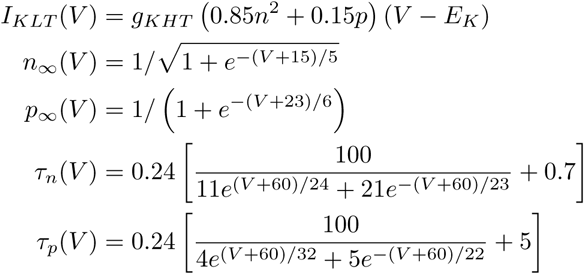

### Hyperpolarization-activated cation current

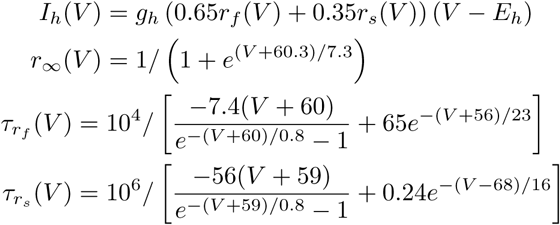

### M-type Potassium current

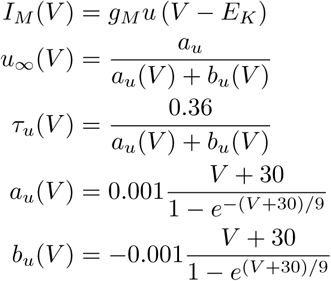

### Sodium current

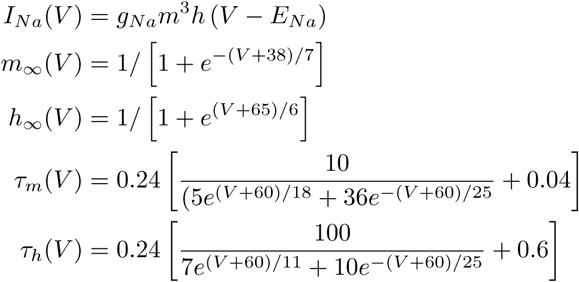

